# Comparing alpha-synuclein-interactomes between multiple systems atrophy and Parkinson’s disease reveals unique and shared pathological features

**DOI:** 10.1101/2024.09.20.613717

**Authors:** S.G. Choi, T. Tittle, R Barot, D. Betts, J. Gallagher, J.H. Kordower, Y. Chu, B.A. Killinger

## Abstract

**Introduction:** Primary synucleinopathies, such as Parkinson’s disease (PD), Dementia with Lewy bodies (DLB), and multiple system atrophy (MSA), are neurodegenerative disorders with some shared clinical and pathological features. Aggregates of alpha-synuclein (αsyn) phosphorylated at serine 129 (PSER129) are the hallmark of synucleinopathies, which for PD/DLB are found predominantly in neurons (Neuronal cytoplasmic inclusions “NCIs”), but for MSA, aggregates are primarily found in oligodendroglia (Glial cytoplasmic inclusions “GCIs”). It remains unclear if the distinct pathological presentation of PD/DLB and MSA are manifestations of distinct or shared pathological processes. We hypothesize that the distinct synucleinopathies MSA and PD/DLB share common molecular features.

**Methods:** Using the in-situ proximity labeling technique biotinylation by antibody recognition (BAR), we compare aggregated αsyn-interactomes (BAR-PSER129) and total αsyn-interactomes (BAR-MJFR1) between MSA (n=5) and PD/DLB (n=10) in forebrain and midbrain structures.

**Results:** For BAR-PSER129 and BAR-MJFR1 captures, αsyn was the most significantly enriched protein in PD/DLB and MSA. In PD/DLB, BAR-PSER129 identified 194 αsyn-aggregate-interacting proteins, while BAR-MJFR1 identified 245 αsyn interacting proteins. In contrast, in the MSA brain, only 38 and 175 proteins were identified for each capture, respectively. When comparing MSA and PD/DLB, a high overlap (59.5%) was observed between BAR-MJFR1 captured proteins, whereas less overlap (14.4%) was observed for BAR-PSER129. Direct comparison between MSA and PD/DLB revealed 79 PD/DLB-associated proteins and only three MSA-associated proteins (CBR1, CRYAB, and GFAP). Pathway enrichment analysis revealed PD/DLB interactions were dominated by vesicle/SNARE-associated pathways, in contrast to MSA, which strongly enriched for metabolic/catabolic, iron, and cellular oxidant detoxification pathways. A subnetwork of cytosolic antioxidant enzymes called peroxiredoxins drove cellular detoxification pathways in MSA. A common network of 26 proteins, including neuronal-specific proteins (e.g., SNYGR3) with HSPA8 at the core, was shared between MSA and DLB/PD. Extracellular exosome pathways were universally enriched regardless of disease or BAR target protein.

**Conclusion:** Synucleinopathies have divergent and convergent αsyn-aggregate interactions, indicating unique and shared pathogenic mechanisms. MSA uniquely involves oxidant detoxification processes in glial cells, while vesicular processes in neurons dominate PD/DLB. Shared interactions, specifically SNYGR3 (i.e., a neuronal protein), between MSA and PD/DLB suggest neuronal axons origin for both diseases. In conclusion, we provide αsyn aggregates protein interaction maps for two distinct synucleinopathies.

## Introduction

Primary synucleinopathies are neurodegenerative diseases, including Parkinson’s disease (PD), dementia with Lewy bodies (DLB), and multiple system atrophy (MSA) that have a common pathological hallmark, specifically alpha-synuclein (αsyn) aggregates^1^. The distribution and abundance of αsyn aggregates are highly heterogeneous between synucleinopathies, with co-pathologies (e.g., tau and beta-amyloid) also being common ^2,3^. MSA is unique among synucleinopathies because αsyn aggregates are predominantly observed in oligodendroglia (glial cytoplasmic inclusions or “GCIs”, also called Pap-Lantos bodies)^4^; in contrast, in the PD/DLB brain neurons bear the majority of Lewy pathology (neuronal inclusions or “NIs”)^1^. NIs are assumed to originate in neurons, but the origins of GCIs are less obvious because, unlike neurons, oligodendroglia do not normally, abundantly, express αsyn^5–7^. Therefore, in the MSA brain, either αsyn aggregates spread from diseased neurons to oligodendroglia or diseased oligodendroglia ectopically express αsyn. There is empirical evidence to support both hypotheses^8^, but details remain unclear.

Research has focused on characterizing NIs in PD/DLB brains, but there is less understanding of the GCIs in the MSA brain, partly due to the lower incidence of MSA (1.6 cases per 100,000 over age of 40) when compared to PD/DLB^9–12^. Recently, we applied an in-situ proximity labeling technique called biotinylation by antibody recognition (BAR) to measure the αsyn interactome in PD/DLB brains and found vesicles, neurons, and neuronal synapses are strongly associated with αsyn aggregates^13^, consistent with the neuronal origin of PD/DLB. BAR has not been applied to other synucleinopathies; in this study, we utilized BAR to compare αsyn interactomes of both diseases to define better core molecular features, distinguishing MSA and PD/DLB.

## Materials and Methods

### Tissue Preparation

Human brain tissues from individuals with a primary clinical diagnosis of MSA, PD, DLB, or Parkinson’s disease dementia (PDD) were obtained from the Rush Movement Disorders Brain Bank (See Table 1 for detailed case information). All brain specimens were prepared following a previously described method^13^. Briefly, 2-cm coronal slabs were fixed by immersing them in a 4% paraformaldehyde solution in 0.1M phosphate-buffered saline pH 7.4 (PBS) for seven days at 4°C. Subsequently, the fixed slabs underwent gradual equilibration in a cryoprotectant solution composed of PBS, 2% dimethyl sulfoxide, and 20% glycerol. Brain regions of interest were cut into 40 μm coronal sections using a freezing stage sliding knife microtome (American Optical). Brain sections were stored in a cryoprotectant solution at -20°C until further processing.

### Immunohistochemistry

Free-floating brain sections were washed in dilution media (“DM,” 5mM Tris-HCl pH 7.6, 150 mM NaCl, 0.05% Triton X100), followed by heat-induced antigen retrieval (HIAR) using sodium citrate buffer (10 mM sodium citrate, 0.05% Tween-20, pH 6.0) at 80°C for 30 minutes. Subsequently, the sections were incubated in a peroxidase quenching solution (0.3% hydrogen peroxide, 0.1% sodium azide) containing blocking buffer (3% goat serum, 2% bovine serum albumin (BSA), 0.4% Triton X-100 in DM) for 1 hour at room temperature. Tissues were then incubated overnight with PSER129 antibody (Abcam, "EP1536Y", ab51253, RRID: AB_869973) or the MJFR1 antibody (Abcam, ab138501, RRID: AB_2537217) diluted to 1:50,000 or 1:20,000 in blocking buffer, respectively. The following day, the sections were washed in DM and incubated with a biotinylated anti-rabbit antibody (Vector Laboratories, BA-1000, RRID: AB_2313606) at a 1:200 dilution in blocking buffer for 1 hour at room temperature, followed by rinsing in DM. Tissues were incubated with an elite avidin-biotin complex (ABC) reagent (Vector Laboratories, PK6100, RRID: AB_2336819) for 75 minutes at room temperature. Tissues were washed in DM and sodium acetate buffer (0.2 M Imidazole, 1.0 M sodium acetate buffer, pH 7.2). Sections were developed using a standard nickel-enhanced 3,3’-diaminobenzidine (DAB)-imidazole protocol, rinsed again with sodium acetate buffer and PBS (50 mM Tris-HCl, pH 7.2, 158 mM NaCl), and then mounted on gelatin-coated glass slides. Counterstaining of the tissues was performed using methyl green. The sections were dehydrated, cleared with xylenes, and coverslipped with Cytoseal 60 (Fisher Scientific). Detailed IHC protocol can be found at protocols.io (https://doi.org/10.17504/protocols.io.8epv5x3mdg1b/v1).

### Microscopy and Imaging

Prepared slides were imaged using an upright microscope (Olympus, BX53) for 4X images. For 20X images and whole section scans, either a Nikon A1 inverted microscope or Odyssey M imager (Li-Cor) was utilized. Images underwent downsizing, cutting, auto-color balancing, and auto-brightness adjustments using Adobe Photoshop (RRID: SCR_014199) to improve clarity in figure and data presentation. The edited images were imported into Adobe Illustrator (RRID:SCR_010279) for arrangement and figure construction.

### BAR

For each case, PSER129 (BAR-PSER129), total αsyn (BAR-MJFR1), and a primary antibody omission negative control (BAR-NEG) were performed essentially as previously described^13^. A notable exception was that, the tyramide reaction was conducted in 100 mL of borate buffer (0.05 M Sodium Borate, pH 8.5) containing 20 μL of stock biotinylated tyramide (Sigma, 5 mg/mL dissolved in DMSO) and 10 μL of hydrogen peroxide (30% H2O2, Sigma-Aldrich) for 30 minutes at room temperature. After BAR labeling, the sections were washed with PBS and placed in a crosslink reversal buffer consisting of 5% SDS, 500 mM Tris-HCl (pH 8.0), 2 mM EDTA, 1 mM PMSF, and 200 mM NaCl. Tissues were heated to 95°C for 30 minutes and incubated at 65°C overnight. The samples were centrifuged to remove insoluble debris, and the supernatant was diluted in TBST (150 mM NaCl, 1% Triton X-100, and 50 mM Tris-HCl, pH 7.6). Biotinylated proteins were captured using 40 μL of streptavidin-coated magnetic beads (Thermo Fisher, Catalog No. PI88817) at room temperature for two hours. The beads were isolated using a magnetic stand (Millipore), washed with excess TBST three times for 30 minutes each, and incubated overnight for 16 hours at 4°C. Following the pulldown, the lysates were retained for later characterization by western blot. Capture proteins were eluted by heating the beads to 98°C in 50 µL of sample buffer containing 1X LDS sample buffer (Invitrogen, Cat. B0008) and 1X sample reducing agent (Invitrogen, Cat. B0009) for 10 minutes. 40 µL of each sample was run on a 4-12% Bis-Tris Gel (Fisher Scientific, Cat. NW04127BOX) for ∼7 minutes at 200 volts (run until sample enters gel). The gel was incubated overnight in a fixation buffer (50% ethanol and 10% acetic acid). The next day, the gel was rehydrated with ultrapure water, stained with Coomassie blue (Invitrogen, LC6060), and the entire lane containing the captured protein was excised for LC-MS/MS analysis.

### LC-MS/MS

Samples were prepared and analyzed by LC-MS/MS using optimized protocols as previously described^14^. Gel pieces containing embedded proteins were digested with trypsin according to optimized protocols. The resulting tryptic peptide mixture was then subject to mass spectrometry analysis.

Determination of BAR-enriched proteins Combining multiple search engines and quantitative software can improve the analysis of MS-based proteomics data sets^3^. Therefore, we combined two common approaches, namely Mascot/Scaffold (i.e., Total normalized spectra, “TNS”) and Andromeda/Maxquant (LFQ). Raw files from each approach were independently processed and analyzed using each approach. Differential abundance (DA) analysis was performed using the R package DEP designed for proteomics analysis^4^. Proteins missing in at least one sample were excluded, data was normalized using variance stabilizing transformation, and the QRILC method imputed missing values. Significant proteins were identified using protein-wise linear models and empirical Bayes statistics using Limma. Proteins from each analysis were combined for downstream pathway analysis. False Discovery Rates (FDR) p-values were adjusted to account for multiple tests, and proteins with q-values below 0.05 were deemed significant. These were estimated using Fdrtool. Proteins significantly enriched for each capture (i.e., significantly enriched over BAR-NEG) were exported and analyzed by multiple list comparator (Molbiotools.com) to determine overlapping and unique proteins identified for each condition. Overlapping proteins were determined for all conditions (i.e., BAR capture, and disease state, Fig. 3C) and individual comparisons within each capture condition (Fig. 4 and 5). Direct comparisons were also made between PD/DLB and MSA for each BAR capture with any known background proteins (i.e., significantly enriched in BAR-NEG) being deemed insignificant.

**Figure. 1.**
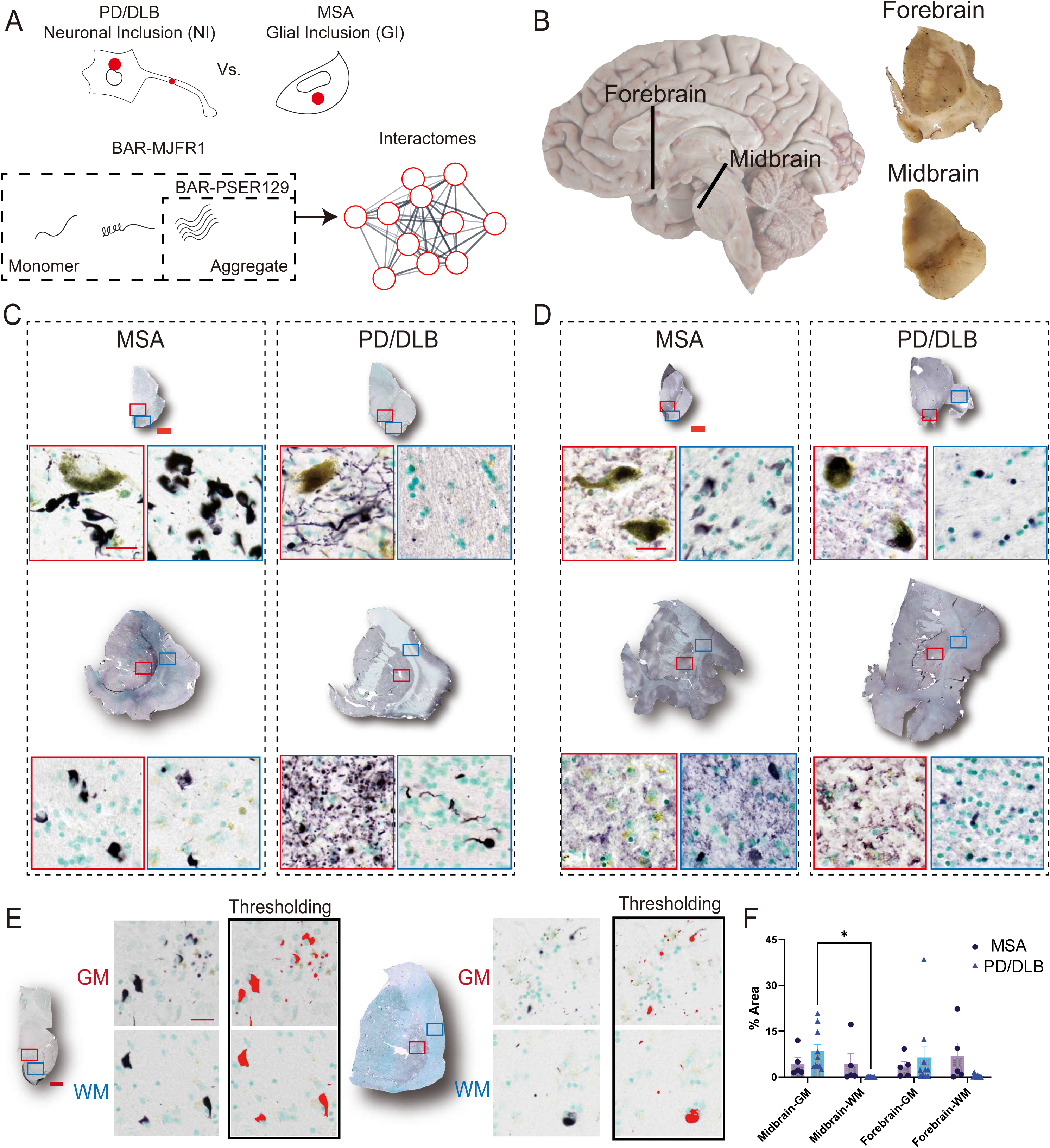
Summary of approach. (A) The distribution of αsyn aggregates is distinct between the synucleinopathies PD/DLB and MSA. For PD/DLB, αsyn aggregates are observed prominently in neuronal cell bodies and projections, termed neuronal inclusions (NI). In contrast, in the MSA brain, αsyn aggregates occur prominently in glia, termed glial inclusions (GI). Biotinylation by antibody recognition (BAR) was used to identify and compare interactomes of total αsyn (BAR-MJFR1) and aggregated αsyn (BAR-PSER129) directly in the PD/DLB and MSA brain. MJFR1 antibody maps to an epitope of a.a. 118-123 of αsyn’s c-terminus and captures physiological monomeric forms as well as aggregates. PSER129 preferentially labels αsyn aggregates, especially in postmortem brain where physiological PSER129 is scarce^58^, and thus BAR-PSER129 will capture aggregate interactions. (B) Depiction of specimen sampling for studies. Longitudinal image of the right hemisphere of the human brain with the approximate location of sampling sites shown (black lines). For BAR, a single coronal section through the Caudate Nucleus and Putamen (Forebrain) and a single transverse section through the midbrain were pooled and used. IHC staining for (C) PSER129 and (D) αsyn (i.e., MJFR1) in forebrain and midbrain sections. Whole-section scans and high-magnification images of select pathology-bearing regions are shown with red and blue boxes denoting the approximate area of the high-magnification image. Sections were stained using nickel- DAB chromogen (black) and counterstained with methyl green (green). Signal thresholding was applied to 20X images of PSER129-stained tissues, specifically in grey matter (GM, red box) and white matter (WM, blue box) in the forebrain and midbrain, for quantification. Enlarged 20X images showing the application of thresholding (E), along with the subsequent quantification of PSER129 immunoreactivity across all cases (F), are presented. Scale bars: C, D = 2 mm and 25 µm; E = 2 mm and 20 µm. MSA, n = 5; PD/DLB, n = 10.

**Figure 2.**
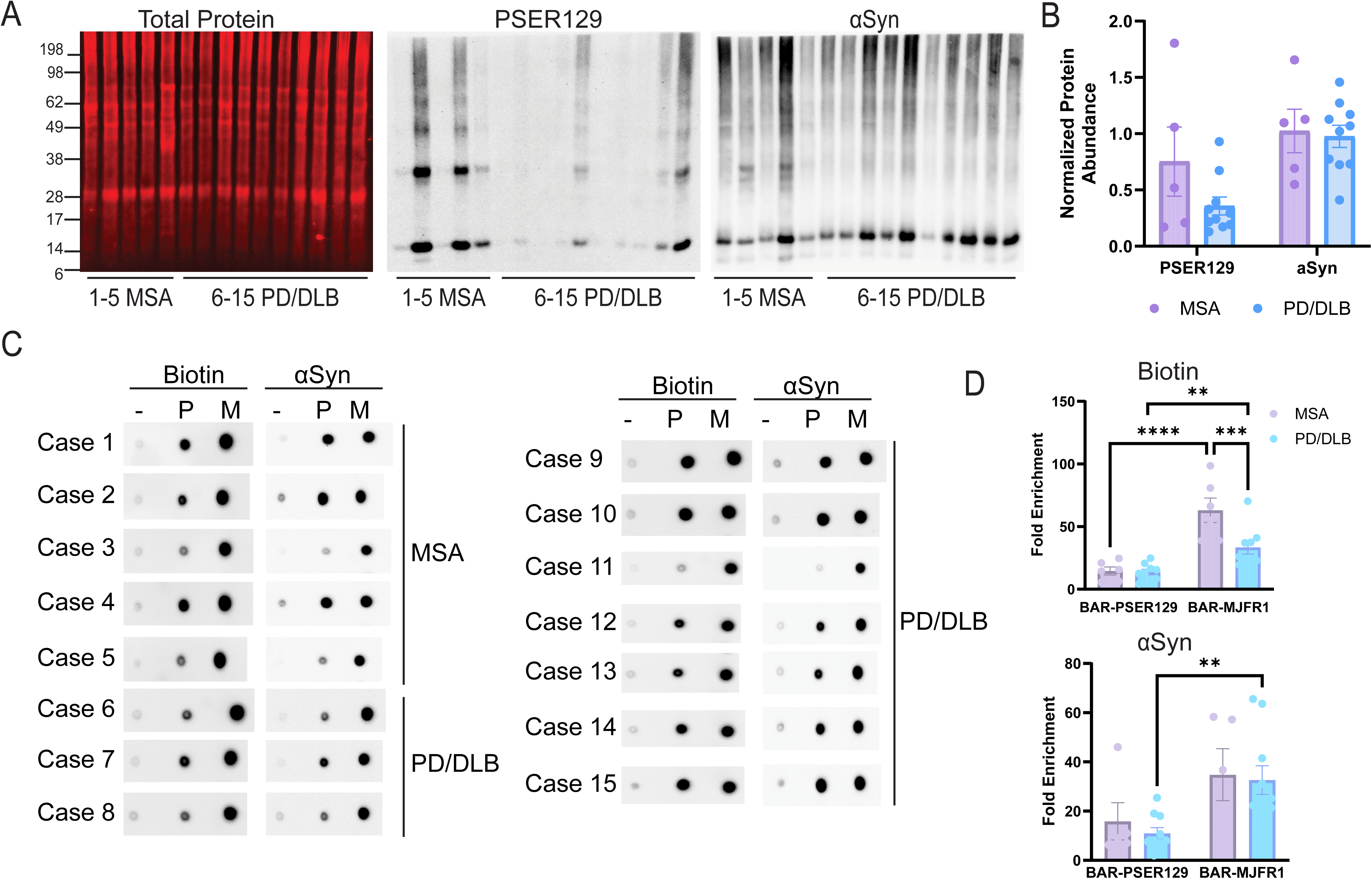
BAR capture in the PD/DLB and MSA brain. (A) 10 μg tissue lysate protein for each case were separated on 4-12% Bis-Tris gel transferred onto PVDF and stained for total protein (Revert total protein stain, LI-Cor), PSER129, or αsyn. Chemiluminescence was used to detect PSER129 and αsyn. (B) PSER129 and αsyn relative density values were first normalized to total protein (i.e., loading control) and then normalized to the mean intensity for each group. (C) Spot blots of 1µl eluent from BAR captures including BAR-NEG (Primary antibody-omission control, “-”), BAR-PSER129 (aggregates “P”), and BAR-MJFR1 (total αsyn “M”) for each synucleinopathy case. Blots were probed for either biotin (ABC reagent) or αsyn (BDSYN1). (D) BAR enrichment was calculated by dividing for “P” and “M” by the relative density value for “-”(i.e., fold-enrichment over background) (mean ± SEM, **P<0.005, ***P<0.005, ****P<0.0001, n=5-10).

**Figure 3.**
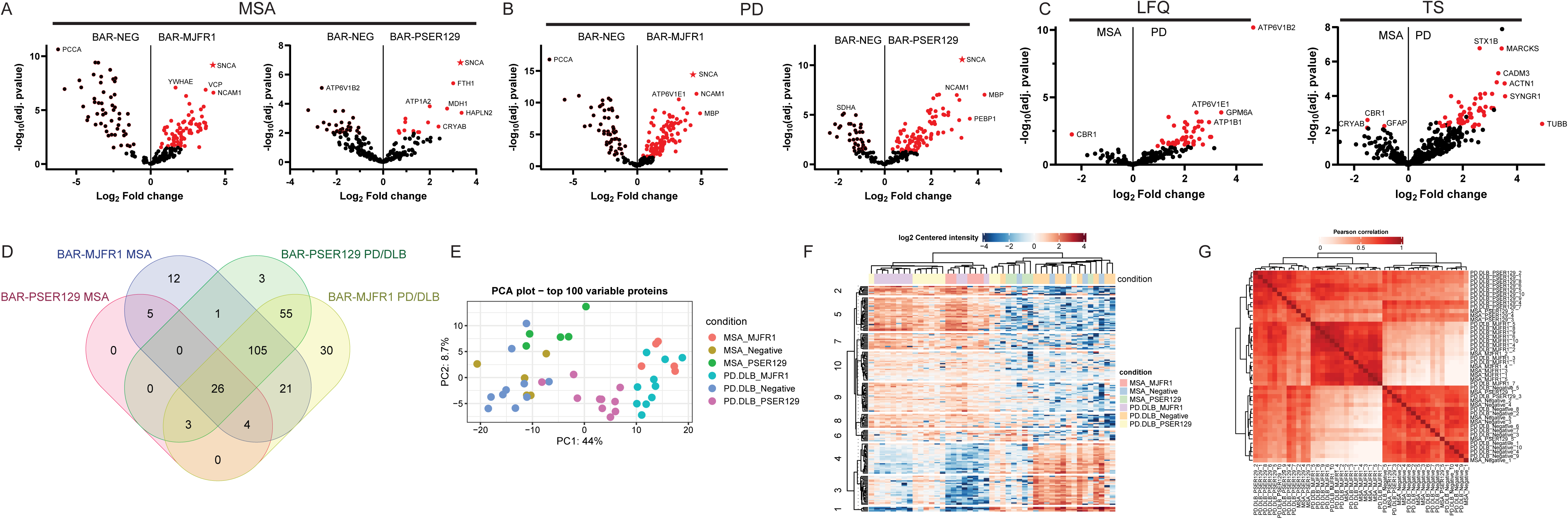
PSER129 and αsyn proximal proteins in the synucleinopathy brain. BAR-labeled proteins were identified by LC-MS/MS and quantified using two approaches Maxquant/andromeda label free quantification (LFQ) and Scaffold/mascot total normalized spectra values (TNS). Volcano plots comparing protein abundance (i.e., LFQ value) between capture (BAR-MJFR1 and BAR-PSER129) and assay background (BAR-NEG) for (A) MSA and (B) PD/DLB. αsyn (i.e., BAR target protein) is denoted as a star. All significant differentially abundant proteins appear red. LFQ results shown, TNS results can be found in Fig. S2. Names of select high abundance proteins are annotated. (C) BAR-PSER129 differentially abundant proteins between MSA and PD/DLB brain. Proteins previously found to be significantly enriched in background (A, B) were excluded for MSA with PD/DLB comparison. Both LFQ and TNS results are shown. (D) Venn diagram showing BAR enriched proteins (i.e., significant over BAR-NEG) for each BAR condition and disease state. Proteins from both analysis (LFQ and TNS) were included. (E) Principal component analysis (PCA) plot of the top 100 variable proteins (See Fig. S9 for TNS PCA plot). (F) Heatmap with non-biased hierarchical clustering of protein abundance across all samples. LFQ shown, for TNS see Fig. S7 (G) Correlation heatmap comparing protein abundances between all samples. LFQ shown, for TNS see Fig. S8. PD/DLB, n=10. MSA, n=5.

**Figure 4.**
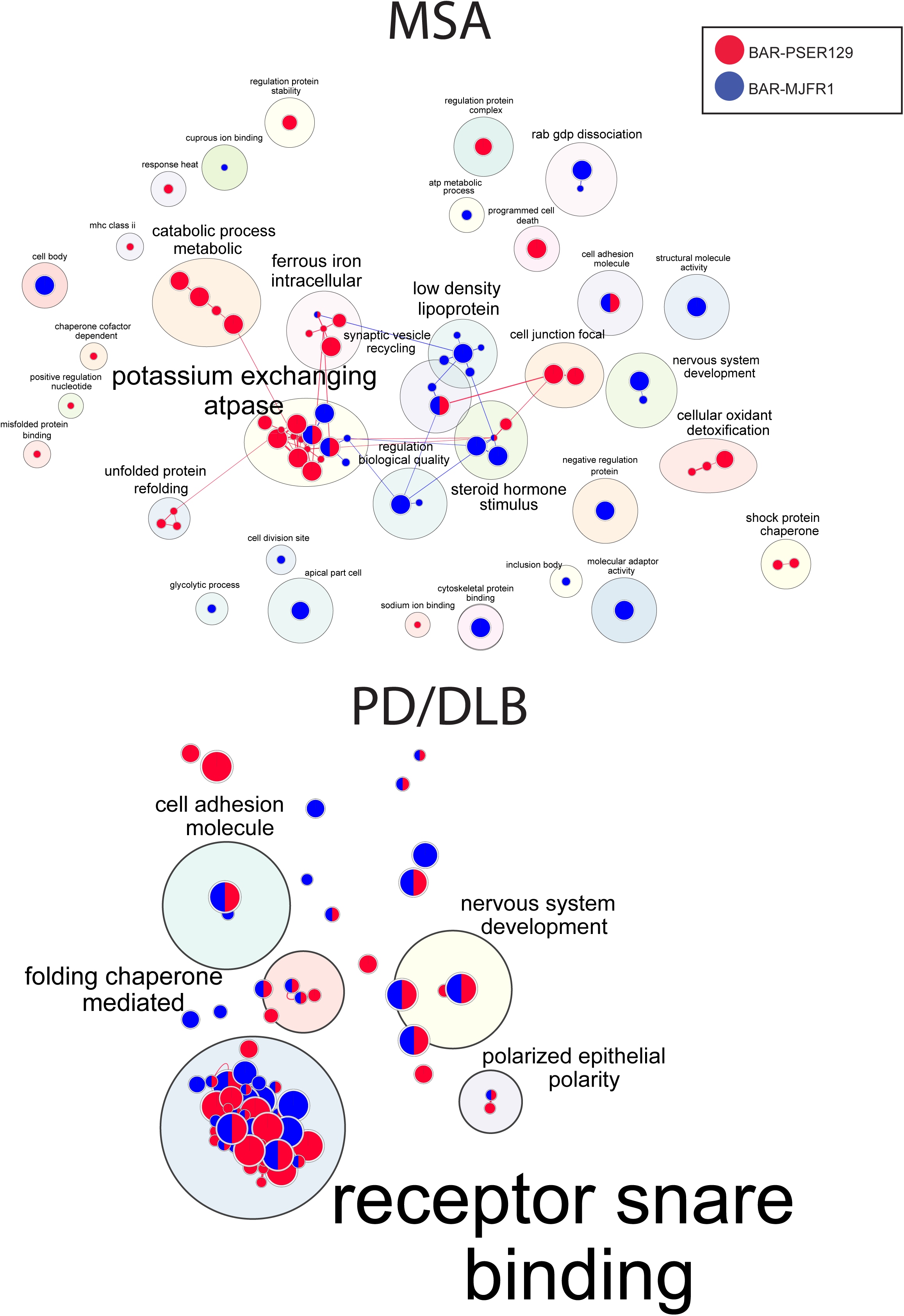
Pathway enrichment map for BAR-identified proteins. BAR-identified proteins from PD/DLB and MSA brains were analyzed by profiler, and significant (q>0.05) GO Driver Terms were mapped using Enrichmentmap and annotated with Autoannotate. GO Driver terms were used to reduce redundancy and simplify enrichment maps. Nodes are color-coded according to BAR capture condition, BAR-PSER129 (Red) or BAR-MJFR1 (Blue). The top-panel is MSA enrichment map, and bottom-panel is PD/DLB enrichment map.

**Figure 5.**
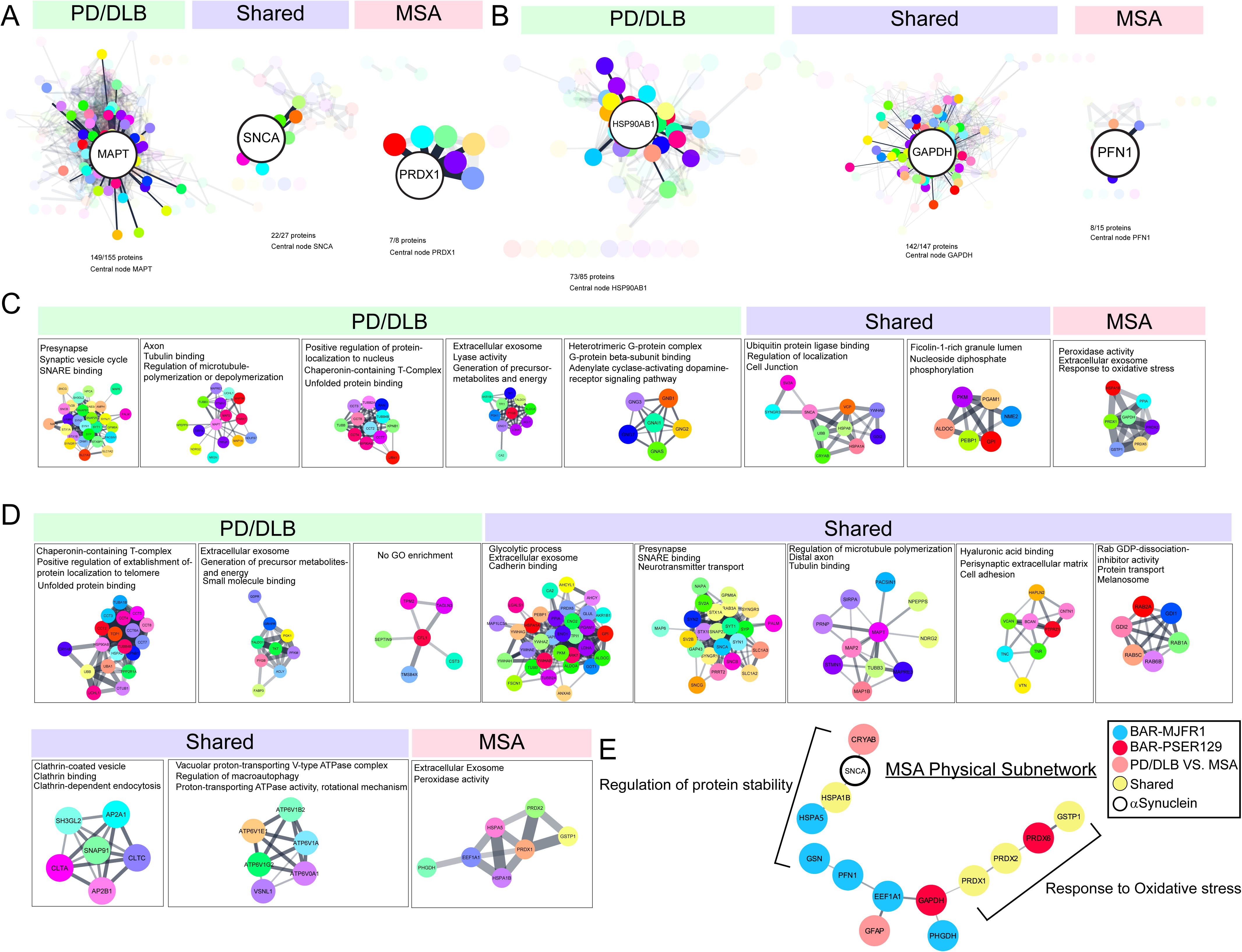
Protein interaction networks identified in PD and MSA brain. BAR identified proteins were analyzed with STRING to plot known functional interactions for each proximal proteome. Proteins were grouped according to whether they were unique to PD/DLB, MSA, or shared between synucleinopathies. STRING networks for (A) BAR-PSER129 and (B) BAR-MJFR1 are shown. CentiScaPe 2.2 was used to determine the central “driver” node for each STRING network (i.e., highest “betweenness” score). The driver node was enlarged, and the first neighbors in network highlighted. For, BAR-PSER129 (C) and BAR-MJFR1 (D) MCL clustering was used to group nodes. Clusters with 6 nodes or more are depicted. Top enrichment (GOCC, GOMF, GOBP) are annotated onto each cluster. (E) Physical interaction network of proteins specific to MSA in for all differential expression analysis.

### Enrichment mapping

Protein lists were input to the gProfiler web user interface. Enrichment analysis was conducted without ranks or weighting because we cannot assume measured protein abundance is a function of distance to BAR target- or relevance. Gene ontology (GO) and Reactome databases were queried without electronic annotations. To reduce redundancy and minimize the size of enrichment map, gProfiler Driver terms exclusively used to construct final enrichment map. Enrichment results were exported to cytoscape and plotted using Enrichmentmap V 3.4.0 with autoannotate.

### STRING

Protein lists were imported in the STRING version 12.0 web user interface. Search was conducted using Homo sapiens as the reference organism. Mapped proteins were then used to generate STRING functional or physical protein-protein interaction networks. The minimum required interaction score was set to 0.5 unless noted otherwise. STRING maps were then exported to Cytoscape version 3.10.2 for MCL clustering, pathway enrichment, network analysis (CentiScaPe 2.2), and network layout configuration.

### Spot blotting

Before conducting LC-MS/MS analysis, 1 µl of streptavidin bead eluent was applied to a methanol-activated polyvinylidene difluoride (PVDF) membrane and fully dried. The dried membrane was then reactivated using methanol, followed by a wash in ultrapure water and post-fixation in 4% paraformaldehyde for 30 minutes. Subsequently, the blots were rinsed with TBST (20 mM Tris-HCl pH 7.6, 150 mM NaCl, 0.1% Tween-20) and immersed in a blocking buffer (TBST supplemented with 5% BSA or 5% dry milk) for 1 hour at room temperature. To detect biotinylated proteins, the blots were treated with prepared ABC reagents (Vector Laboratories) in the blocking buffer for 30 minutes at room temperature. For αsyn detection, the blots were incubated overnight with anti-αsyn antibody "SYN1" (BD Lifesciences, #610787, RRID:AB_398108) diluted 1:2000 in the blocking buffer. After the incubation, the blots were washed and incubated with anti-mouse HRP conjugate (Cell Signaling, CST7076S, RRID:AB_10956588), 1:6000 in the blocking buffer, respectively. Following another round of washing with TBST, all membranes were imaged using enhanced chemiluminescence (ECL) substrates (Bio-Rad, product # 170–5060) and a Chemidoc imager (Bio-Rad).

### Western blotting

10 µg of protein from the bead eluent were loaded on 4 to 12% gradient Bis-Tris gels (ThermoFisher) and run until loading dye reached bottom of gel. Resolved proteins were blotted onto the PVDF membrane using wet transfer at settings of 100 V for 1.5 hours. Following the transfer, the membranes were rinsed in ultrapure water (18Ω) and then fixed with 4% paraformaldehyde for 30 minutes at room temperature. Membranes were dried completely and reactivated with methanol. The gels were then stained with Revert Total Proteins Stains (Li-Cor, product # 926-11016) according to the manufacturer’s protocol and imaged using an Odyssey M imager (Li-Cor). The membranes were then rinsed in TBST (20 mM Tris-HCl pH 7.6, 150 mM NaCl, 0.1% Tween-20) and blocked in TBST containing 5% BSA or dry milk for 1 hour at room temperature. Primary antibody incubation was done with SYN1 (BD Lifesciences, dil. 1:2000) or PSER129 (Abcam, dil. 1:50,000) diluted in the blocking buffer overnight at 4 °C. The next day, membranes were washed in TBST and incubated for 1 hour with either anti-rabbit (dil. 1:20,000) or anti-mouse (dil. 1:6,000) HRP-conjugated secondary antibodies (Cell Signaling Technologies) diluted in the blocking buffer. The membranes were washed and developed using a chemiluminescence (ECL) substrate (Product no. 1705060, Bio-Rad) with a Chemidoc imager (Bio-Rad). For quantitative blots, chemiluminescent blots were developed using Bio-Rad “optimal auto-exposure” which constructs a quantitative image.

### Quantitative Analysis of Blotting Results

The mean abundance of PSER129 or αsyn for western blot and spot blots was quantified using ImageJ (version 1.54 h, https://imagej.net/ij/download.html, RRID:SCR_003070). The data was then normalized to the total protein amount, or fold enrichment was calculated based on negative controls. Statistical analysis and graphing were performed using GraphPad Prism (Version 10.2.0, https://www.graphpad.com/, RRID: SCR_002798). One-way ANOVA with Tukey’s post-hoc test or Two-tailed unpaired t-test was used to compare experimental groups.

### Quantification of IHC

Brightfield images were captured using an inverted confocal microscope equipped with a 20X objective (Nikon A1R). Annotation of each tissue section was performed within a bounding box of 2000×2000 pixels. All images underwent auto-exposure and auto white balance adjustments using NIS-Elements software. For quantification, RGB-based color thresholding was initially fine-tuned to measure dark black pixels (i.e., PSER129 signal). This adjusted threshold was recorded using the Macro function in NIS-Elements and then uniformly applied in batch to all images, irrespective of the disease (MSA or PD/DLB) or the region of the brain (WM or GM in the midbrain or forebrain) (See supplementary figure 1 for the raw images and how the thresholding was applied). Subsequently, the percentage area of the thresholded signals was exported and graphed using GraphPad Prism. The statistical analysis was performed using Two-way ANOVA with Tukey’s multiple comparisons test.

## Results

To compare MSA with PD/DLB, we performed in situ proximity labeling techniques BAR-MJFR1 and BAR-PSER129^13^, which capture total αsyn and aggregated αsyn spatially (∼50-100nm radius) associated proteins, respectively (Approach summarized in Fig. 1A). BAR-identified proteins were used to describe interactomes and compare the molecular characteristics of these two distinct synucleinopathies. To perform all studies, we used 5 clinically diagnosed MSA and 10 clinically diagnosed PD/DLB cases (See Table 1 for case details). Brain hemisections of the nigrostriatal pathway were used for all studies (Fig. 1B) because both brain regions are involved with the core clinical motor features of both MSA and PD/DLB, and the nigrostriatal pathway bears αsyn aggregates for these synucleinopathies. Tissue sections were positionally matched for all cases and consisted of the forebrain regions (e.g., putamen, caudate, adjacent cortical regions) and the midbrain (e.g.,substantia nigra).

### αSyn in the MSA and PD/DLB forebrain and midbrain

Tissues were first characterized by immunohistochemistry (IHC). Striatal containing forebrain and midbrain hemisections were stained for PSER129 and total αsyn (MJFR1). Results show widespread PSER129 pathology throughout the forebrain and midbrain sections (Fig. 1C). For PD/DLB cases, PSER129 pathology was dense in grey matter with substantia nigra, caudate, putamen, and the insular lobe was often also severely affected. In contrast, MSA cases had widespread PSER129 pathology in the white matter tracts of the midbrain and forebrain, consistent with known white matter involvement for MSA^15,16^ (Fig. 1C). The morphology of PSER129-positive structures in the PD/DLB and MSA brains was consistent with prominent NIs or GCIs^1,4^, respectively. MSA and PD/DLB exhibited αsyn (MJFR1) reactivity throughout the forebrain and midbrain sections, predominantly in the grey matter (Fig. 1D). GCIs were evident in the white matter because αsyn is of low abundance in white matter tracks of the healthy mammalian brain^17^. This makes αsyn bearing oligodendroglia visible, even at low magnification and with total αsyn staining. In contrast, the NIs of PD/DLB observed with PSER129 staining were obscured by a strong ubiquitous total αsyn signal throughout the grey matter (Fig. 1D). Despite the apparent regional specificity of PSER129 staining for MSA and PD/DLB within some cases, a quantitative comparison of PSER129 between white and grey matter revealed high heterogeneity between cases and few overall significant differences in PSER129 abundance between MSA and PD/DLB (Fig. 1E, F). For PSER129 abundance in PD/DLB midbrain, significant differences (Tukey’s post-hoc test p<0.05) were observed between grey and white matter (Fig 1F). Collectively, these results highlight the presentation of αsyn pathology and distribution of BAR targets for the cases used for these studies.

### BAR capture in MSA and PD/DLB forebrain and midbrain tissues

After verifying the distribution and morphology of the αsyn pathology consistent with PD/DLB and MSA, BAR was performed as previously described^18^ to capture proteins within proximity (predominantly within ∼50-100nm radius^19^) to PSER129 (i.e., αsyn aggregates) and αsyn. For each case, a positionally matched forebrain and midbrain section were pooled (Fig S2 average 81.7± 0.021 mg combined wet mass). Western blot (WB) of proteins extracted from pooled forebrain and midbrain hemisections revealed that PSER129 and αsyn abundance was variable between cases (Fig. 2A, B), in agreement with IHC (Fig.-1F); however, PSER129 or αsyn abundance was not significantly different between PD/DLB and MSA (Fig. 2B, PSER129; Two-tailed unpaired t-test, t(13)=1.656, p=0.1215; αsyn Two-tailed unpaired t-test, t(13)=0.2505,p=0.8061). High molecular weight species (>17kDa) were observed on WB for PSER129 and αsyn, which may be SDS-resistant aggregates^20^ or residual formalin crosslinks^21^. αSyn truncation/cleavage products were also observed for most samples, which have been widely reported in healthy and diseased tissues^20,22^.

Following BAR, dot blots of BAR-captured proteins (i.e., bead eluent) revealed biotin and αsyn enrichment were similar between BAR captures for MSA and PD/DLB brains (Fig 2C, D). Biotin enrichment was significantly more for BAR-MJFR1 when compared to BAR-PSER129, regardless of disease state (Fig. 2D, Two-way ANOVA, F (1, 13) = 47.68, P<0.0001). Biotin content was higher for BAR-MJFR1 capture in the MSA brain compared to PD/DLB (Fig. 2D, Sidak’s multiple comparison, p=0.0006). For PD/DLB, αsyn enrichment was significantly more in BAR-MJFR1 than BAR-PSER129 (Fig. 2D,

Sidak’s multiple comparison, p=0.0046). Together these results demonstrate successful BAR-PSER129 and BAR-MJFR1 enrichment for all cases in this cohort.

### BAR-identified αsyn and PSER129 interactomes in the MSA and PD/DLB brain

After filtering missing proteins, we quantified 440 (TNS method) and 209 (LFQ-method) proteins. In the MSA brain, we found 38 BAR-PSER129 enriched proteins and 175 BAR-MJFR1 enriched proteins over the background (i.e., BAR-NEG). In the PD/DLB brain, we identified 194 BAR-PSER129 enriched proteins and 245 BAR-MJFR1 significantly enriched proteins. The BAR target αsyn (SNCA) was the most enriched protein for all captures when quantified by LFQ-method (Fig. 3A and 3B). For the TNS method, SNCA was enriched over the background but was not the most abundantly enriched capture protein (Fig. S3). BAR-NEG enrichment was observed for all contrasts, and the known background biotin-binding protein propionyl-CoA carboxylase subunit alpha (PCCA)^23^ was often most abundant in BAR-NEG (Fig. 3A, B). For BAR-MJRF1, direct comparison between MSA and PD/DLB did not reveal any differentially abundant proteins. In contrast, for BAR-PSER129 direct comparisons revealed numerous proteins more abundant in PD/DLB including ATP6V1B2, STX1B and MARCKS (Fig. 3C). Four proteins (CBR1, GFAP, CRYAB) were differentially abundant in MSA (Fig. 3C). Overlap of identified proteins was greatest (105 proteins, 39.6% of all proteins) between groups excluding BAR-PSER129 in the MSA brain (Fig. 3D). Remarkably, despite the prominent GCI abundance observed in our MSA samples, not a single unique protein was identified for BAR-PSER129 in the MSA brain (Fig. 3D). However, BAR-MJFR1 and BAR-PSER129 identified proteins unique to MSA including HSPA1B, PRDX1, SERTIN8, GSTP1, and PRDX2. Identifying unique proteins was minimal in the MSA brain, and most proteins overlapped with PD/DLB (BAR-MJFR1 and BAR-PSER129). The principal component analysis (PCA) plot of the top 100 BAR-captured proteins showed the separation of BAR-MJFR1 from BAR-NEG in both MSA and PD/DLB brains, with close grouping observed within each disease state (Fig. 3E). Similar to BAR-NEG, BAR-PSER129 in MSA brain was more related to PC2 than PC1.

Fig. 3F and 3G heatmaps show the hierarchical clustering of protein abundance (LFQ) and hierarchical clustering of Pearson correlations between all tested samples, respectively. Cases formed 2 main clusters (Fig. 3F), with one cluster having a strong “on-target” signal, containing the majority of BAR-MJFR1 captures and BAR-PSER129 captures in the PD/DLB brain. The second main cluster was comprised of background samples (BAR-NEG for MSA and PD/DLB), as well as BAR-PSER129 captures in MSA and PD/DLB brains showing weaker “on-target” signals. In terms of correlations, BAR-MJFR1 and BAR-NEG were closely correlated between MSA and PD/DLB, while BAR-PSER129 was correlated less between MSA and PD/DLB.

### Enrichment maps for MSA and PD/DLB

Next, we performed pathway enrichment analysis and identified hundreds of significantly enriched pathways (p-adj.<0.05) for each experimental group (see Supplementary Data for enrichment files). To reduce the redundancy of identified pathways, we used the gProfiler “go driver term’s” function, which first groups significant terms by GO relation and then uses a simple greedy search strategy to identify leading gene/protein sets. The resulting top 10 driver terms are shown in the supplementary Fig S4, and all driver terms were used to build the enrichment map in Figure 4. For both MSA and PD/DLB, most clusters contained pathways enriched for both BAR captures, however unique pathway enrichment was observed for BAR-PSER129. For MSA, BAR-PSER129 unique major clusters included “catabolic/metabolic processes”, “unfolded protein refolding”, “cellular oxidant detoxification”, “cell junction focal”, and “shock protein chaperone” clusters. For PD/DLB, all major clusters identified contained BAR-MJFR1 enriched pathways, and therefore, no wholly BAR-PSER129 specific enrichments were detected. “Receptor/SNARE binding” was the major cluster for PD/DLB.

### Protein interaction networks identified in the synucleinopathy brain

To better understand proteins and pathways that distinguish MSA from PD/DLB, STRING was used to plot functional and physical interactions between BAR identified proteins. Each STING networks’ central node (i.e., driver protein) was determined using CentiScaPe 2.2 and highlighted in Fig. 5A, B. Driver proteins for BAR-PSER129 included MAPT (i.e., tau), SNCA, and PRDX1 for PD/DLB, shared, and MSA, respectively. Driver proteins for BAR-MJFR1 included HSP90AB1, GAPDH, and PFN1 for PD/DLB, shared, and MSA, respectively. MCL clustering was conducted to isolate functional networks for each experimental group. Results showed that for PD/DLB BAR-PSER129 identified mostly neuron-specific networks, including SNARE, presynapse, and axon (Fig. 5C), in agreement with our previous observation (Fig. 4). Shared networks for BAR-PSER129, which included SNCA, involved ubiquitin ligase binding, localization, and follin-1-rich granule lumen. A single network of 7 interconnected nodes enriched for peroxidase activity, extracellular exosome, and response to oxidative stress was unique to BAR-PSER129-MSA. For BAR-MJFR1 (5D), MSA specific network was also associated with extracellular exosome and peroxidase activity. Physical protein interactions for MSA-proteins were plotted in relation to BAR-target protein (i.e., SNCA). An MSA specific physical subnetwork emerged with periodoxin’s (PRDX1, PRDX2, PRDX6) as major components. SNCA was segregated from the MSA specific periodoxin network, with CRYAB, HSPA1B, and HSPA5 being connected to SNCA. This BAR identified subnetwork of MSA was cytosolic (e.g., GAPDH, GFAP), with regulation of protein stability and response to oxidative stress being the major functional themes.

## Discussion

Here, we interrogated αsyn interactomes within the human brain of two clinically and pathologically distinct primary synucleinopathies. We found total αsyn interactions (i.e., BAR-MJFR1) were overall similar between MSA and PD/DLB brains with strong enrichment for presynaptic vesicle processes, broadly consistent with physiological αsyn interactions, αsyn functionality, and previous proximity proteomics studies^13,14,24–26^. In contrast, αsyn aggregate interactions (i.e., BAR-PSER129) were markedly different between MSA and PD/DLB, with several glial-enhanced proteins (CBR1, CRYAB, and GFAP) (Human Protein Atlas proteinatlas.org^27^)^28^ being more abundant in MSA αsyn aggregates (i.e., PD vs. MSA contrast). In addition, five proteins were wholly unique to MSA, including HSPA1B, PRDX1, PRDX2, SEPTIN8, and GSTP1. Collectively identified MSA proteins are functionally interconnected and are involved with anti-oxidative mechanisms and protein stability (Fig. 5E). Multiple oxidative stress mechanisms have been implicated in MSA, including oxidative phosphorylation dysfunction and reductions in anti-oxidative machinery^12^. Reduction in antioxidant glutathione (GSTP1 identified here) may occur in MSA^29,30^, but peroxiredoxins or CBR1 (both proteins protect from oxidative stress) have not been directly implicated in MSA. Together, our findings implicate unique anti-oxidative mechanisms for MSA pathology. We cannot infer precisely how these proteins fit the MSA pathological process, whether protective or toxic. Interestingly, reductions in antioxidant COQ10 have been implicated in MSA pathogenesis^31–33^, albeit inconsistently^34^, and clinical efficacy trials for ubiquinol are currently underway^35^.

Our data implicates canonical αsyn pathways (i.e., vesicles/SNARE) for PD/DLB and anti-oxidative pathways for MSA but also demonstrates shared interactions between the two distinct synucleinopathies. All experimental conditions shared 26 proteins involved in cellular functions such as intracellular iron sequestering, protein refolding, and nucleoside diphosphate phosphorylation, representing the core interactome of αsyn for both diseases (Fig. S10). At the center of this network was HSPA8, a regulator chaperonin-mediated autophagy inhibitor of αsyn aggregation, implicated in PD, DLB, MSA, and other neurodegenerative diseases^36,37^. Therefore, our data suggests a protein-refolding response at the center of disparate synucleinopathies. Despite the predominating GCIs in MSA, several neuronal vesicle proteins, including SYNGR3 and SV2A, were identified in the shared pathway (Fig. S10), supporting a neuronal origins hypothesis for MSA. Both proteins could originate in the few PSER129 reactive neuropils in our MSA cases (Fig.1), or they might originate in vesicles released from neurons and endocytosed by proximal oligodendrocytes ^8,38–40^. Indeed, extracellular exosomes (i.e., L1CAM positive^41,42^) were consistently enriched pathway in agreement with our previous BAR studies^13^ suggesting αsyn aggregates in synucleinopathy brain partly resemble exosomes, and supporting the hypothesis that αsyn/αsyn aggregates can spread via exosomes^43–46^. It’s unclear if αsyn’s “exosomeness” has biological significance (i.e., αsyn aggregates are being spread by exosomes) but αsyn containing exosomes have been isolated from peripheral sources^47,48^. Alternatively, coincidental exosome enrichment may occur because of αsyn/αsyn aggregates close association with vesicles/endosomes. Our data suggests one of two scenarios, αsyn aggregates are spreading to oligodendrocytes via presynaptic neuronal exosomes or in addition to αsyn several neuronal proteins are being upregulated (SYNGR3 and SV2A) in diseased oligodendrocytes, but this is unlikely because differentiating oligodendrocytes do not express these genes^28^. Overall, our findings support a hypothesis for the potential late-stage contribution of oligodendrocytes to the disease process for MSA^8,49^.

We identified CRYAB associated with GCI’s. CRYAB is upregulated in the MSA brain and directly associated with GCIs^50^. Recent studies have identified GCI-like structures in oligodendrocytes of PD patients carrying αsyn gene mutations or presenting atypical clinical features^51^. CRYAB is encoded by a subpopulation of oligodendrocytes that is depleted in sporadic PD^52^. Functionally, CRYAB binds to the ends of αsyn fibrils, inhibiting aggregation and potentially playing a protective role. The strong association of CRYAB with GCI-related diseases, such as MSA, suggests that this “capping” mechanism plays a prominent role and may contribute to the particularly aggressive clinical progression observed in MSA. Additionally, MSA-derived seeds extracted from the human brain exhibit significantly higher templated-mediated seeding efficiency, which may explain the pronounced CRYAB upregulation and its sequestration to αsyn inclusions. These findings support the potential therapeutic utility of small heat shock proteins in managing these neurodegenerative conditions.

The study’s limitations include that most of the MSA cases here are MSA-Parkinsonian type (MSA-P), which is the dominant subtype of MSA in the western hemisphere^53–55^, restricting the comparison of PD/DLB to only one subtype of MSA.

Another limitation is BAR as a “Spatial-omics”^56^ technique has special interpretation considerations, including spatial resolution (∼100nm radius from BAR target), lack of information about precise nature of protein-protein interactions (i.e., direct or indirect), and although cell type/cell compartment can be inferred from BAR data it cannot be determined. Another limitation is our analysis focused on MSA, networks/proteins specific for PD/DLB identified here are likely valuable for follow-up analysis deciphering PD/DLB mechanisms.

Proximity proteomics are powerful techniques used to study αsyn aggregate and non-aggregate interactomes^13^, but this is the first application of BAR comparing synucleinopathies and there currently aren’t standardized data analysis pipeline specifically for BAR. As we found before^13^, background captures (BAR-NEG) were a critical component to the study design that allowed identification of BAR-antibody specific proteins. Direct comparisons between disease (e.g., MSA vs PD/DLB) could be made for each BAR capture, but background proteins must be removed manually, as background (BAR-NEG) information is lost in this approach. To increase protein identification and robustness of our analysis we conducted two analyses based on two quantitative values LFQ scores (Maxquant) and TNS (Scaffold). We found correlation between methods (Fig S5,6) but LFQ was superior in that the target protein (αsyn) was consistently the most enriched protein, in contrast to TNS where αsyn was only most abundant in the MSA-PSER129 capture (Fig. S3).

## Conclusion

Our results suggest that MSA involves PRDX’s anti-oxidative processes in glial cells, while presynaptic vesicle/SNARE processes dominate PD/DLB. Our findings revealed a significant overlap between MSA and PD/DLB, suggesting shared pathogenic origins for these synucleinopathies. Intriguingly, we identified several neuronal proteins, besides SNCA, and strong exosome enrichment, associated with GCIs, together supporting the neuron-to-glia spread hypothesis of MSA.

## Data availability

The mass spectrometry proteomics data have been deposited to the ProteomeXchange Consortium via the PRIDE^57^ partner repository with the dataset identifier PXD05530

## Supporting information

Protein lists

MSA MJFR1 gprofiler_DRIVER

MSA MJFR1 gprofiler_All

MSA PSER129 DRIVER

MSA PSER129 All

PDDLB MJFR1 All

PDDLB MJFR1 DRIVER

PDDLB PSER129 DRIVER

PDDLB PSER129 All

## Acknowledgments

Data processing and statistical analysis were performed partly by VUGENE (Vilnius, Lithuania). Valuable bioinformatics support was also provided by Miglė Gabrielaitė (VUGENE). Proteomics services were performed by the Northwestern Proteomics Core Facility, generously supported by NCI CCSG P30 CA060553 awarded to the Robert H Lurie Comprehensive Cancer Center, instrumentation award (S10OD025194) from NIH Office of the Director, and the National Resource for Translational and Developmental Proteomics supported by P41 GM108569. Work was supported by Rush Movement Disorders Pilot grant via a generous donation by the Postuma family. National Institutes of Health 5R01NS128467 also supported work.

## Notes

### Competing Interest Statement

The authors have declared no competing interest.

