## Supplemental Appendix for "Comparing alpha-synuclein-interactomes between multiple systems atrophy and Parkinson’s disease reveals unique and shared pathological features"

**Table of Contents:**

Figure S1- Thresholding on midbrain and forebrain images
Figure S2- Tissue weights used for experiments.
Figure S3- Volcano plots for TNS method.
Figure S4 – Top ten Driver pathways.
Figure S5 – Comparison of LFQ and TNS methods for BAR-PSER129.
Figure S6 - Comparison of LFQ and TNS methods for BAR-MJFR1.
Figure S7- Heatmap for TNS method.
Figure S8- Correlation heatmap for TNS method.
Figure S9- PCA plot for TNS method.
Figure S10-STRING network for 26 proteins common to all captures and disease states.

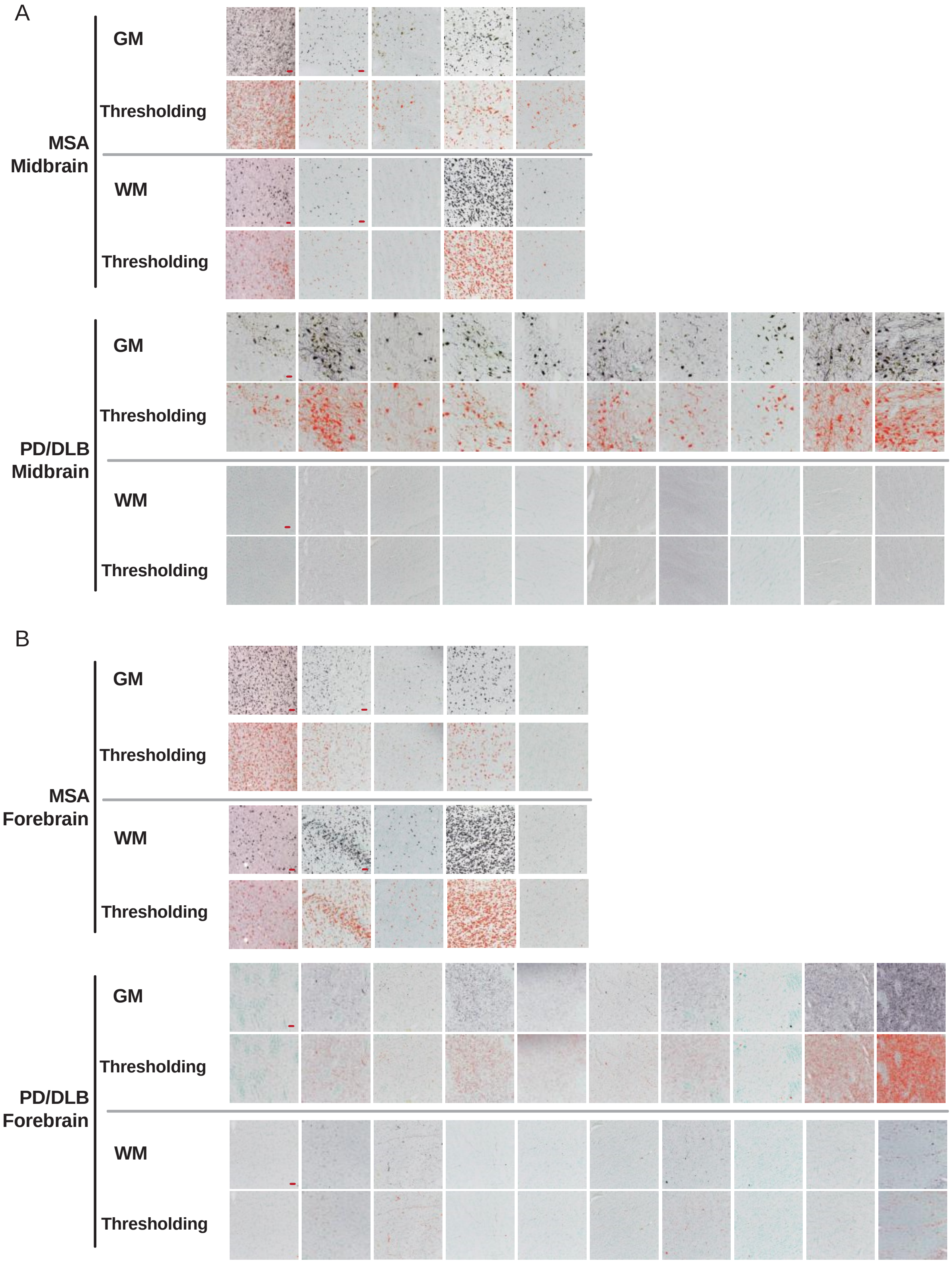


**Figure S1**. Thresholding applied to midbrain and forebrain images. To quantify the PSER129 signals in each region, 20x images of (A) midbrain and (B) forebrain from MSA and PD/DLB were batch-applied with thresholding. In the midbrain, GM images were taken at the SNpc, while WM images were captured at the white matter tract near the GM. For the forebrain, GM images were taken at the putamen, and WM images were taken at the white matter tract near the putamen. Scale bar = 50 μm.

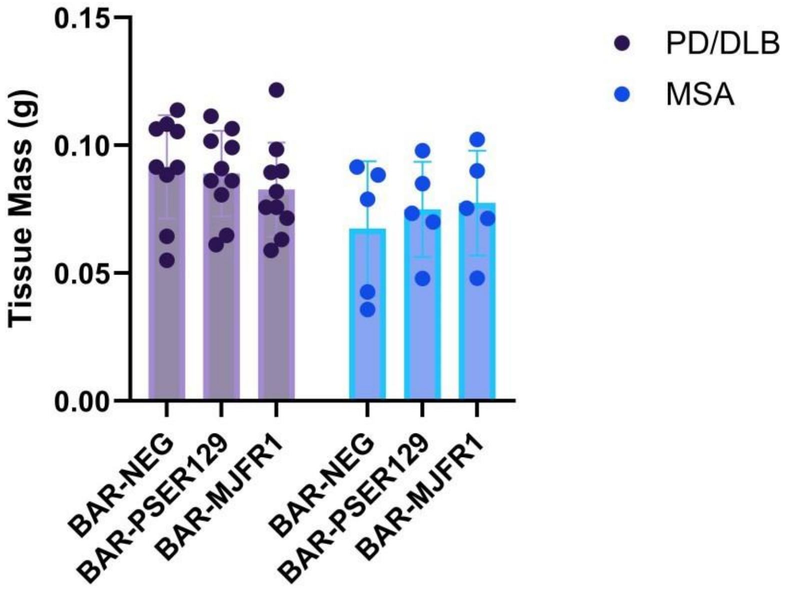


**Figure S2**. Tissue weight input for BAR. A single forebrain and midbrain section were pooled for each BAR capture. Prior to processing for BAR, wet tissues were weighed for each capture. Graph shows the tissue weights for all BAR samples prepared. PD/DLB, n=10 and MSA, n= 5.

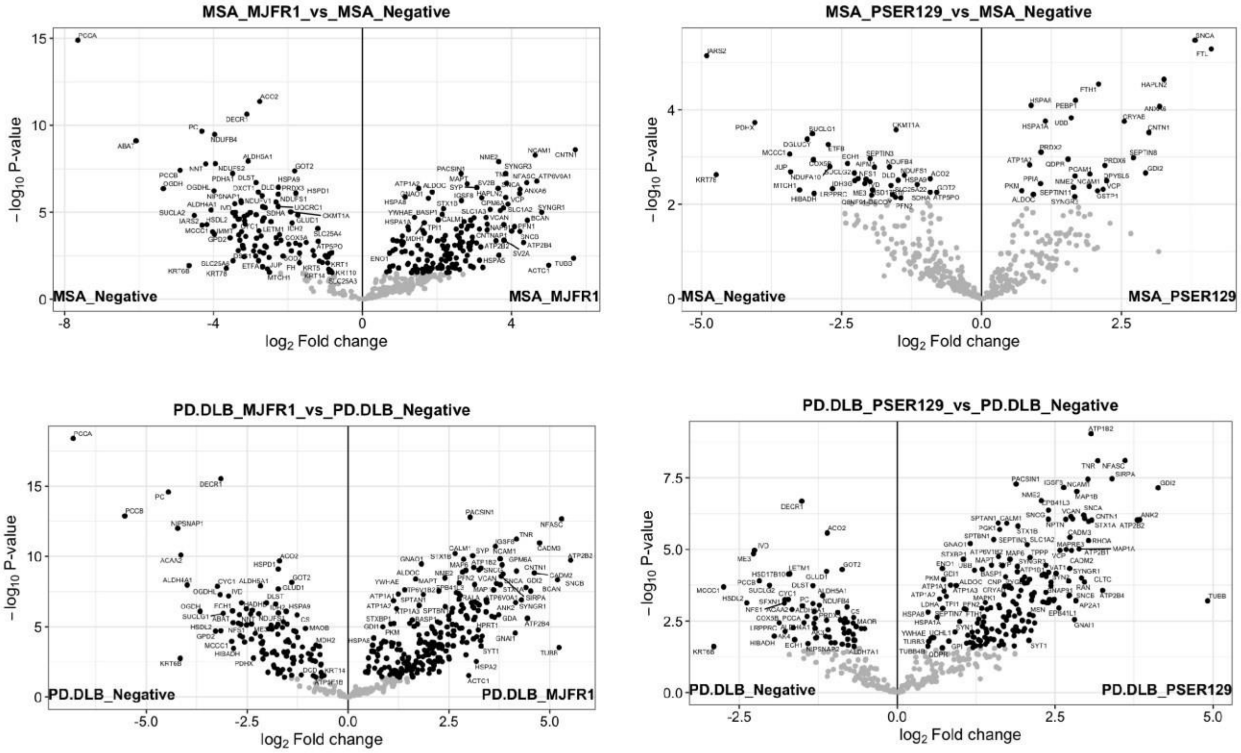


**Figure S3.** Volcano plots for TNS method. Plots display the proteins quantified for each condition using the TNS method. For BAR-PSER129 in MSA, SNCA was the most enriched protein, while other captures included SNCA above the background but did not rank it as the top hit.

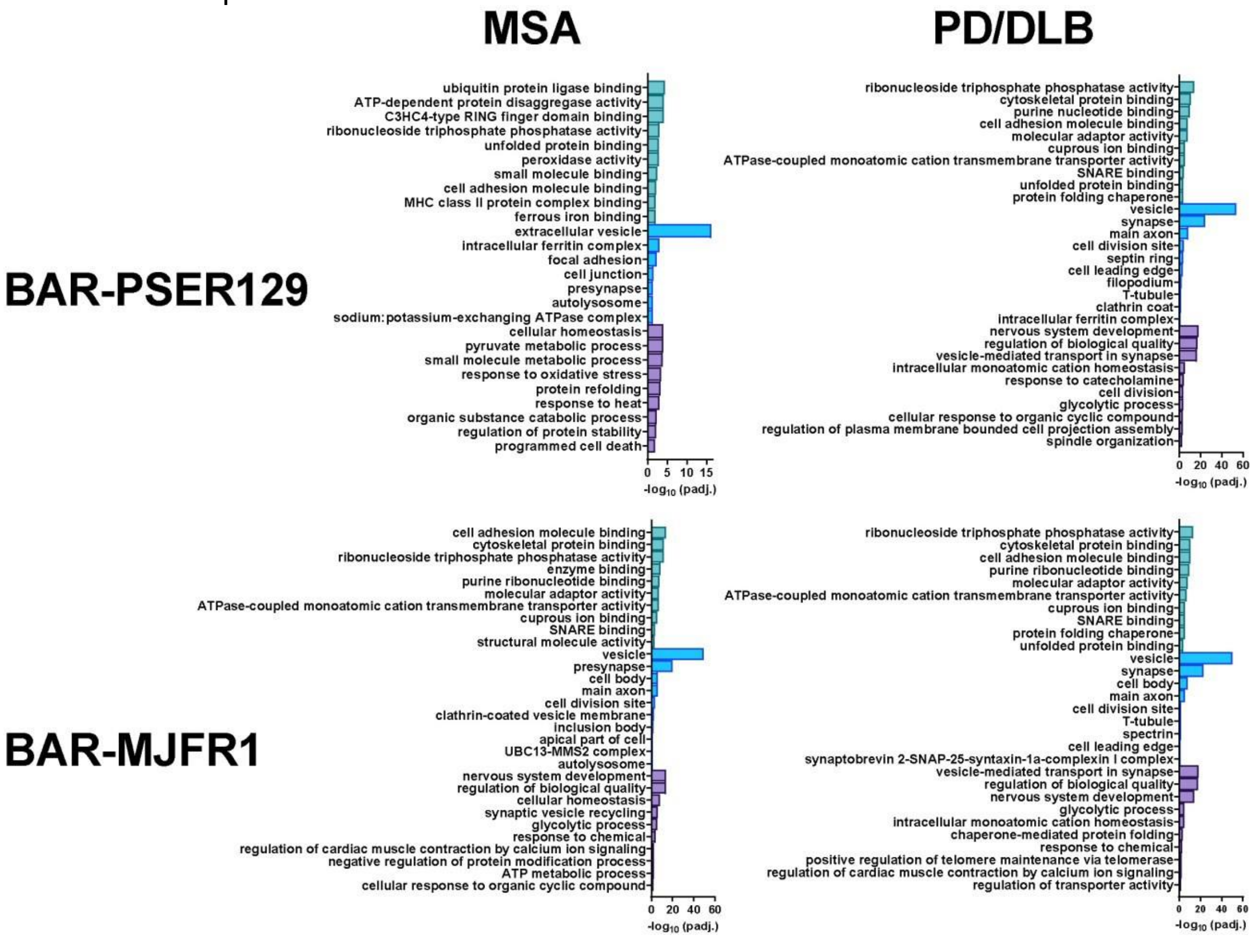

**Figure S4. Top-10 enriched GO pathways.** gProfiler enrichment was conducted on BAR-identified proteins. Graphs show the top 10 (ranked by padj.) driver GO pathways for each condition. All driver GO pathways are found in the enrichment map, Figure 4.

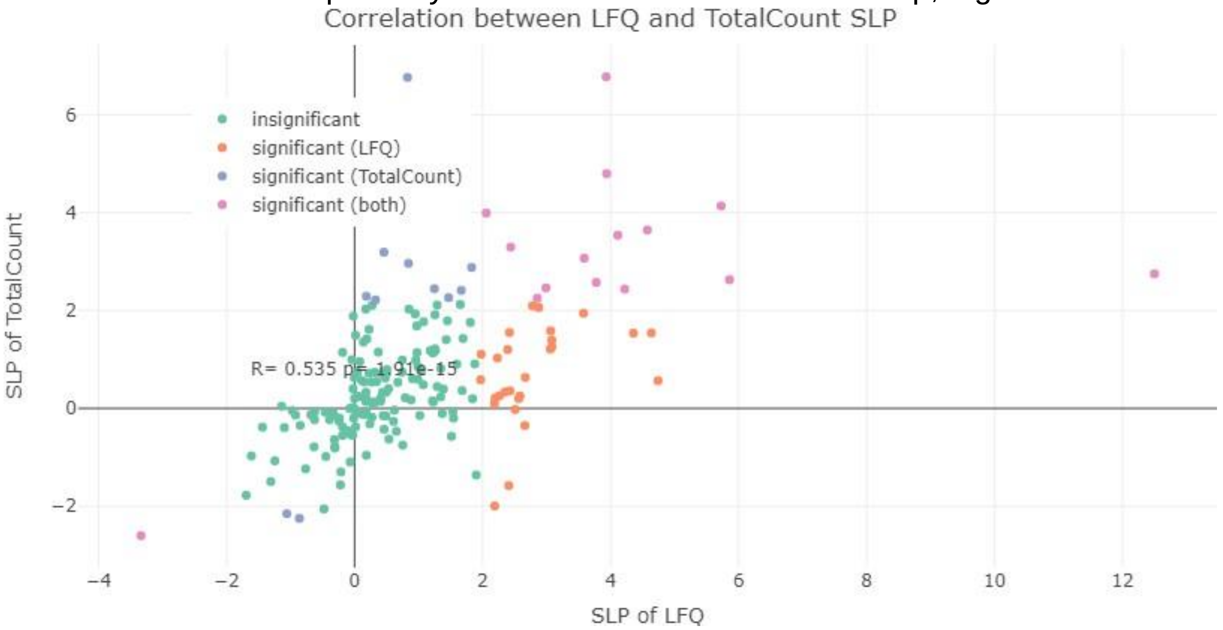


**Figure S5**. Comparison of LFQ and TNS ("total count") methods for BAR-PSER129. Sine log-p-value (SLP) was calculated for differential abundance between LFQ and TNS methods. This method takes p-value and log fold-change into account. Only proteins captured in both analyses are included in the comparison. Pearson R was calculated and displayed on the graph.

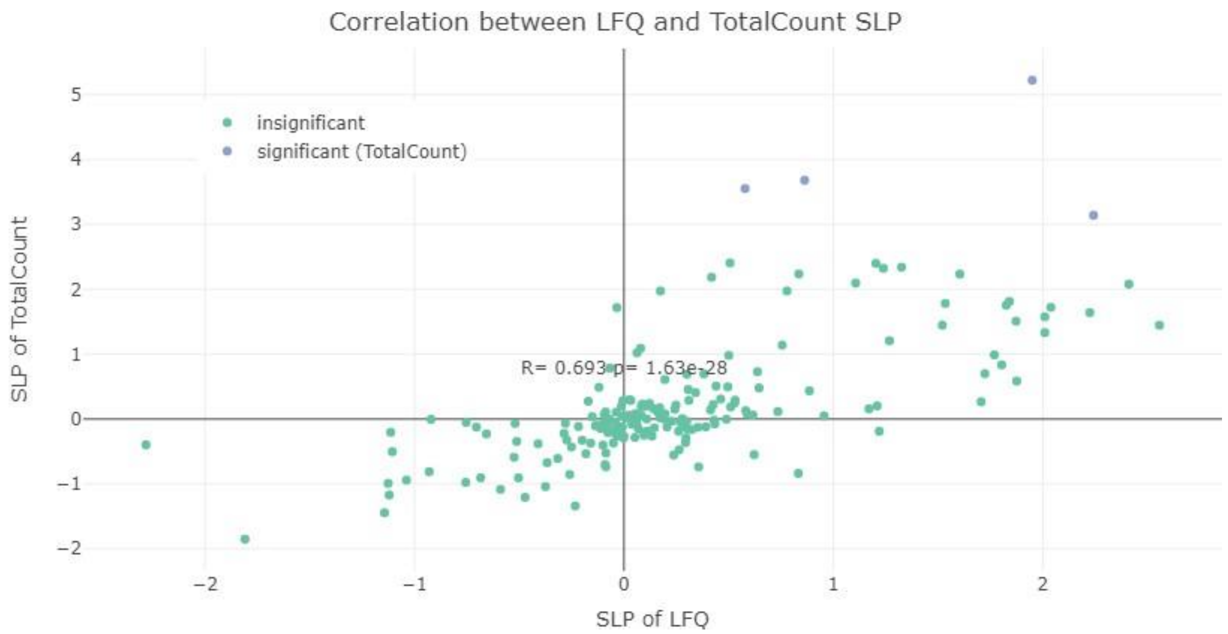


**Figure S6**. Comparison of LFQ and TNS (“total count”) methods for BAR-MJFR1. Sine log-p-value (SLP) was calculated for differential abundance between LFQ and TNS methods. This method takes p-value and log fold-change into account. Only proteins captured in both analyses are included in the comparison. Pearson R was calculated and displayed on the graph.

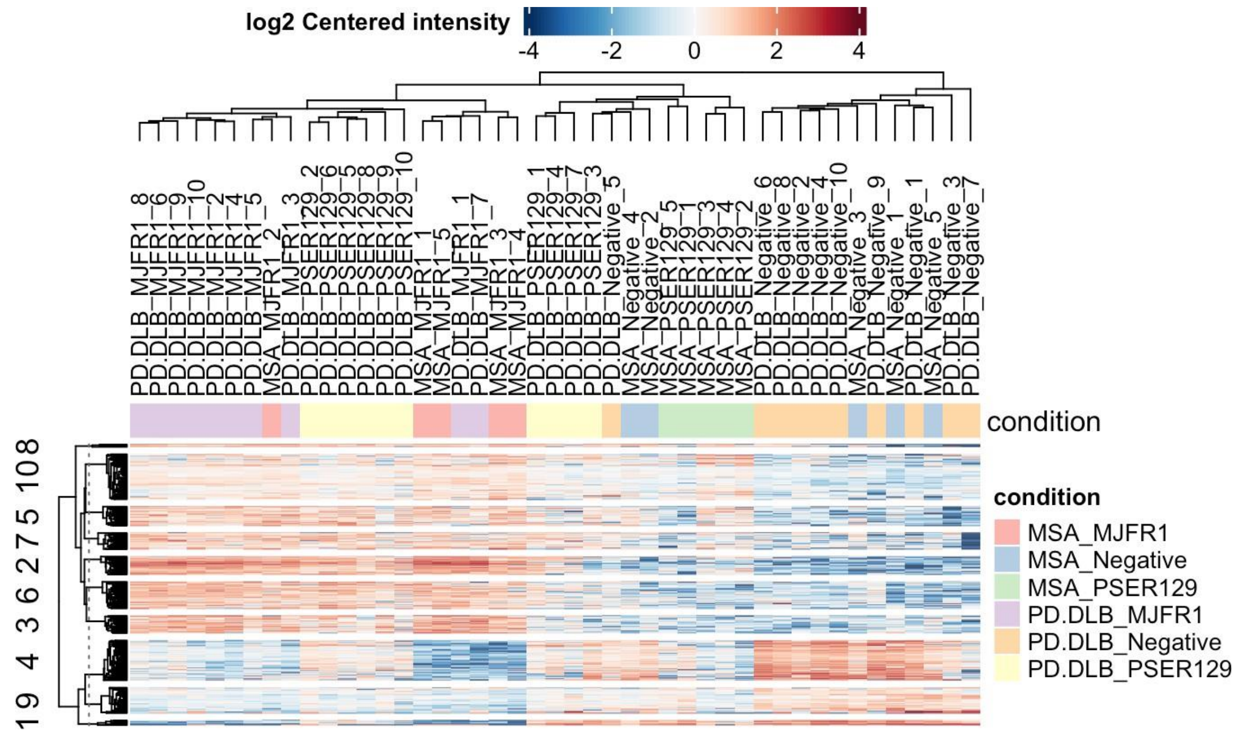


**Figure S7**. Heatmap for TNS method. Hierarchical clustering of protein abundance among all tested samples. The cases formed two main clusters: one cluster included BAR-Neg for both MSA and PD/DLB, while the other main cluster showed strong “on-target” signals for BAR-MJFR1 in MSA and PD/DLB, as well as for BAR-PSER129 in PD/DLB. Although BAR-PSER129 in MSA was included in this cluster, it exhibited weaker “on-target” signals.

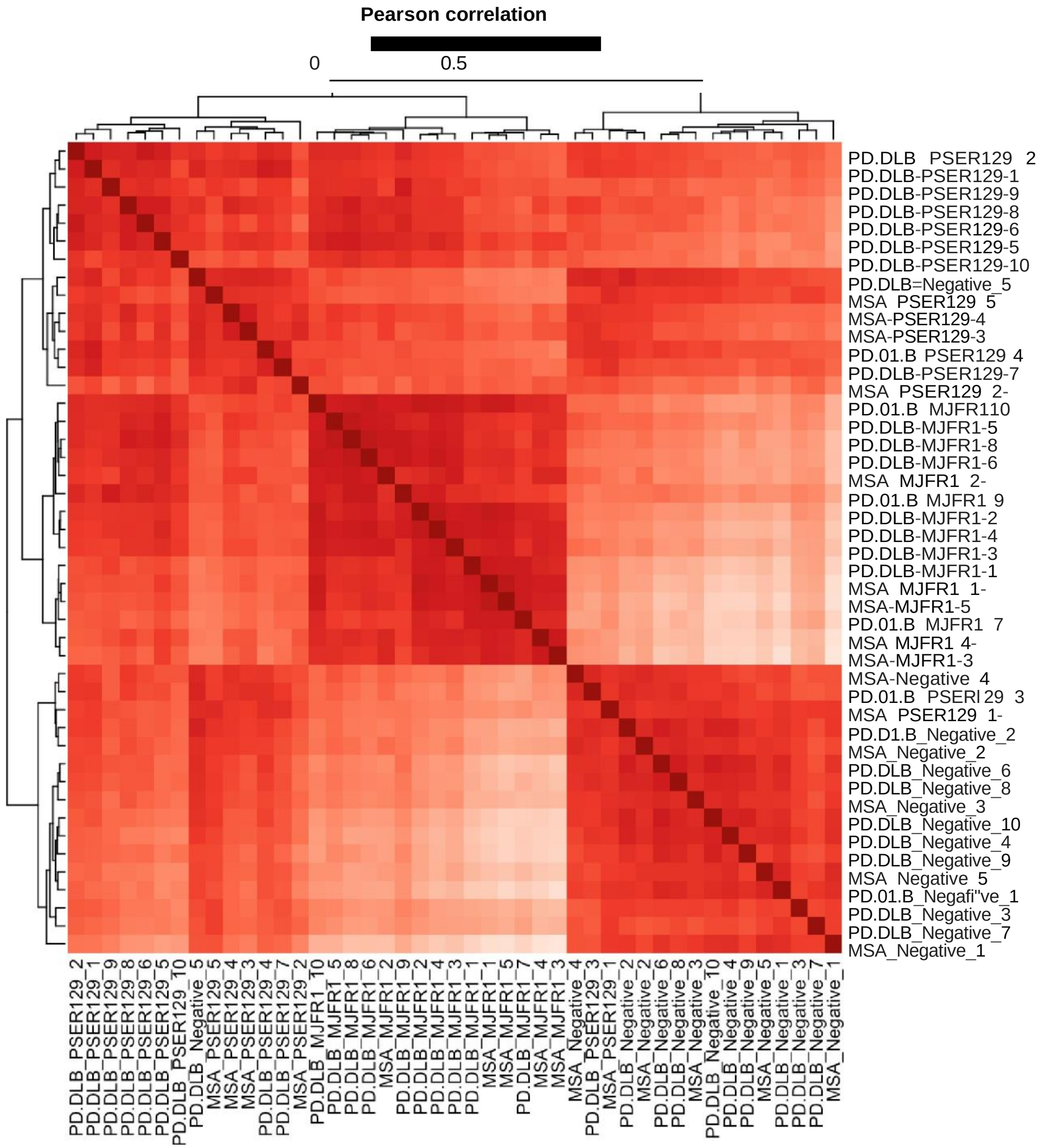


**Figure S8**. Correlation heatmap for TNS method. Hierarchical clustering of Pearson correlations among all tested samples. BAR-MJFR1 showed a close correlation between MSA and PD/DLB, as did BAR-Neg between the diseases.

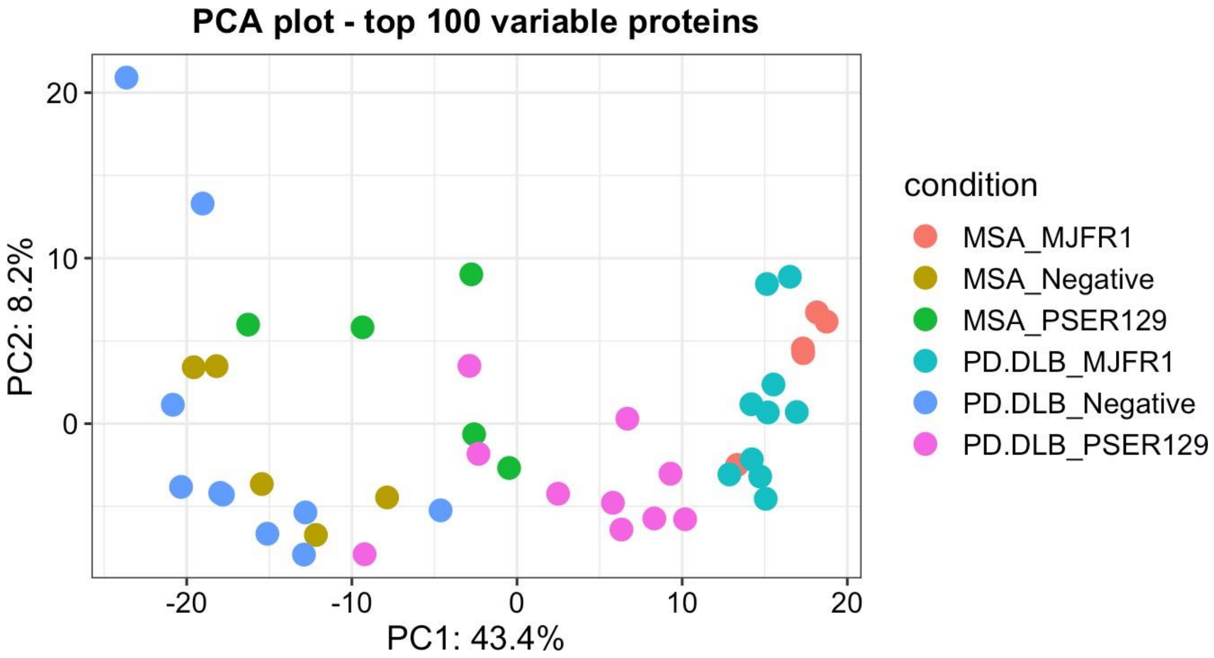


**Figure S9**. PCA plot for TNS method. PCA plot for the TNS method, which displays the top 100 BAR-captured proteins, reveals that BAR-MJFR1 in MSA and PD/DLB exhibit close groupings, overlapping in both PC1 and PC2. BAR-PSER129 for MSA is positioned near the BAR-Neg captures, while the captures in PD/DLB are close to the BAR-MJFR1 captures.

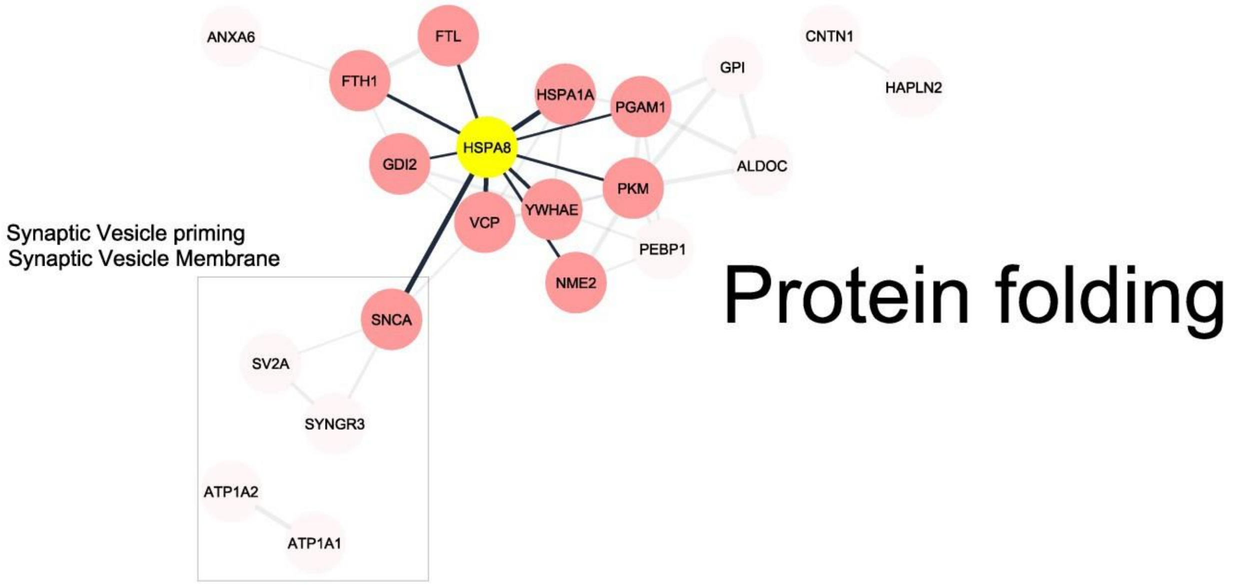


**Figure S10**. STRING network for 26 proteins common to all captures and disease states. 26 proteins were identified and enriched for all captures and disease states. STRING network of these proteins reveals a network with HSPA8 as the central node and pathways involving protein folding as the major enrichments. Interestingly, several presynaptic proteins (SV2A and SYNGR3) are included in this network, despite the lack of neuronal pathology observed in MSA.
