## Supplementary material for "Comparing alpha-synuclein-interactomes between multiple systems atrophy and Parkinson’s disease reveals unique and shared pathological features": Table 1

**Table 1.** *Characteristics of MSA and PD/DLB patients.*

| **Sample #** | **Sex** | **Age at death (yrs)** | **Method of fixation** | **Hoehn and Yahr Scale** | ***Disease Duration (yrs)** | **Clinical diagnosis** | **Pathological diagnosis** |
| --- | --- | --- | --- | --- | --- | --- | --- |
| 1 | M | 64 | Immersion fixation 7-10 days (same for all) | NA | NA | MSA | MSA-C |
| 2 | M | 55 | Immersion fixation 7-10 days (same for all) | NA | NA | MSA | MSA-P |
| 3 | F | 75 | Immersion fixation 7-10 days (same for all) | 4 | 4 | MSA-P | MSA-P |
| 4 | F | 65 | Immersion fixation 7-10 days (same for all) | 5 | 10 | MSA-P | NA |
| 5 | M | 60 | Immersion fixation 7-10 days (same for all) | 3 | 2 | MSA-P | NA |
| 6 | M | 74 | Immersion fixation 7-10 days (same for all) | NA | 9 | PD | Corticobasal degeneration with parkinsonism. Alzheimers |
| 7 | F | 87 | Immersion fixation 7-10 days (same for all) | 3 | 2 | PD | LBD. Primary age related tauopathy |
| 8 | M | 81 | Immersion fixation 7-10 days (same for all) | 4 | 3 | PD | PD-brainstem prominent |
| 9 | M | 84 | Immersion fixation 7-10 days (same for all) | 4 | 4 | PD | NA |
| 10 | M | 80 | Immersion fixation 7-10 days (same for all) | 4 | 6 | DLB | DLB diffuse subtype |
| 11 | m | 79 | Immersion fixation 7-10 days (same for all) | 4 | 8 | PD | NA |
| 12 | F | 89 | Immersion fixation 7-10 days (same for all) | 2 | 4 | PD | NA |
| 13 | M | 80 | Immersion fixation 7-10 days (same for all) | 4 | 8 | PD | NA |
| 14 | M | 73 | Immersion fixation 7-10 days (same for all) | 2 | 11 | LBD | NA |
| 15 | M | 70 | Immersion fixation 7-10 days (same for all) | 4 | 5 | PD/PSP | NA |

* The duration of the disease is defined as the period from the onset of symptoms to death, as documented in the clinical records of the Rush Movement Disorder program.
